# SNPD-siKRAS: siRNA specifically inhibits KRAS with a single-nucleotide mutation

**DOI:** 10.1101/2025.09.18.677211

**Authors:** Yoshiaki Kobayashi, Yoshimasa Asano, Atsushi Sato, Hiroaki Taniguchi, Kumiko Ui-Tei

## Abstract

*Kirsten Rat Sarcoma Viral Oncogene Homolog* (*KRAS*) is the most frequently mutated gene associated with pancreatic cancer. However, targeting mutated *KRAS* without affecting the wild-type KRAS expression by conventional chemotherapy is challenging, because *KRAS* protein is considered to be undruggable due to the lack of surface structures for drug binding. Here, we developed a single-nucleotide polymorphism-distinguishable small interfering RNA (SNPD-siRNA) that suppresses the expression of mutated *KRAS* without affecting the wild type *KRAS* expression through discrimination of single-nucleotide differences using RNA interference technology. In two- and three-dimensional cultures of human pancreatic cancer-derived cell lines, SNPD-siKRAS significantly reduced the mutated *KRAS* expression, and suppressed the cell proliferation by inhibiting the activity of MAPK/PI3K pathway. Furthermore, the xenograft experiments using mice revealed that the growth of implanted pancreatic cancer-derived cells were suppressed by the SNPD-siKRAS. Thus, this technology may provide a therapeutic platform for personalized genomic medicine that can identify the single nucleotide mutations in the disease-causing genes.

## INTRODUCTION

*Kirsten Rat Sarcoma Viral Oncogene Homolog* (*KRAS*) is an oncogenic driver gene in various cancers. A total of 46,821 single-nucleotide mutations of the *KRAS* gene in cancers are registered in the Catalogue Of Somatic Mutations In Cancer (COSMIC, v. 96)^1^. Among those mutations, 45,999 are located in codons 12 and 13 within the guanosine triphosphate (GTP)-binding domain of KRAS (Figures 1A and 1B). The wild-type (WT) *KRAS* gene is a molecular switch, shifting between the GTP-bound active form and guanosine diphosphate (GDP)-bound inactive form^2–4^ (Figure 1C). GTP binding to KRAS plays a critical role in cellular proliferation, differentiation, and death^5^ through the activation of multiple downstream effector pathways, including the mitogen-activated protein kinase (MAPK) and phosphatidylinositol 3-kinase (PI3K) pathways^6,7^. KRAS mutations in codons 12 and 13 are non-synonymous mutations and mutated (Mut) KRAS accumulates in an active form, disrupting hydrolysis from GTP to GDP (Figure 1C) and thereby promoting continuous downstream signaling for tumor cell proliferation^8,9^. These mutations are present in 98% of pancreatic cancers, the highest level among cancer types, and in 45% and 31% of colorectal and lung cancers, respectively. Over the past 40 years, despite great efforts to develop drugs targeting Mut KRAS, targeting KRAS proteins with conventional drugs has remained challenging because KRAS does not have a suitable surface structure for binding such drugs. One small-molecule inhibitor related to a KRAS (G12C) mutation, sotorasib, was recently approved as the first covalent inhibitor of Mut KRAS^10^. However, KRAS (G12C) represents only 11% of all KRAS mutations^11^, and therefore the applicability of this drug is limited. No other direct KRAS therapies have yet been established; most existing therapies are indirect, focusing on the inhibition of the downstream signaling pathway of KRAS. Therefore, a more universal therapeutic strategy is needed.

**Figure 1.**
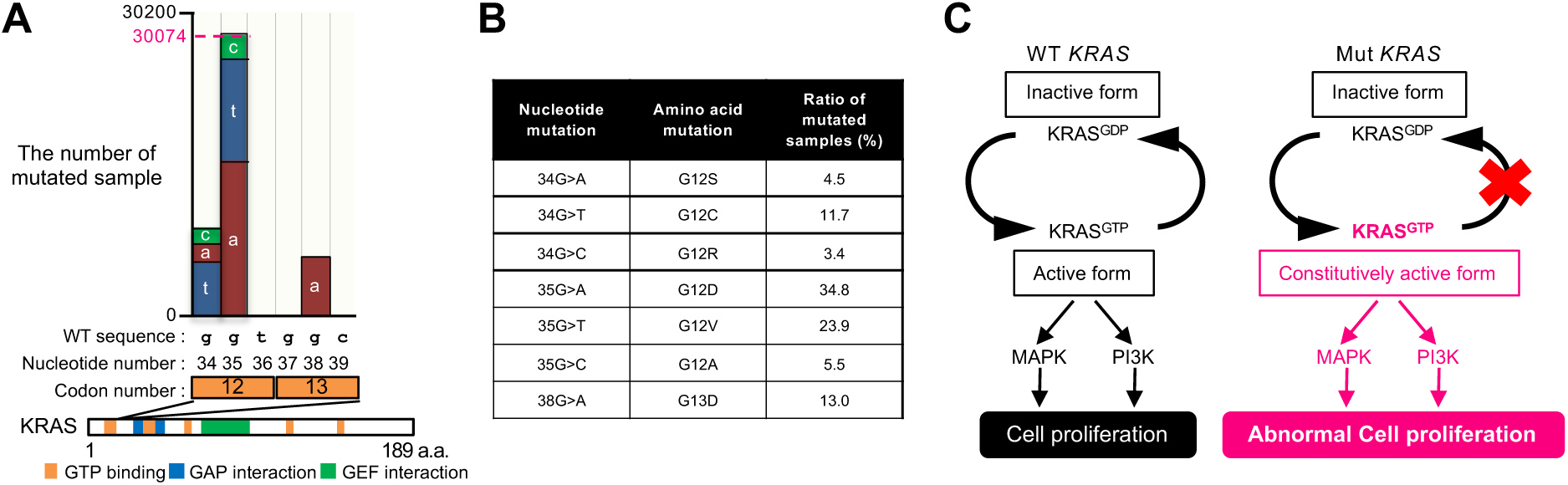
Distribution of single-nucleotide mutations in KRAS. (A) Distribution of single-nucleotide mutations in codons 12 and 13 of KRAS. The lower panel shows the domain structure of KRAS, consisting of 189 amino acids (a.a.). The numbers of mutations were 9,179 at nucleotide position 34 and 30,074 at nucleotide 35 in codon 12, as well as 589 at nucleotide 37 and 6,157 at nucleotide 38 in codon 13. (B) List of single-nucleotide mutations and corresponding amino acid mutations with mutation ratios > 1% in codons 12 and 13. (C) Schematic presentation of G protein-binding state of KRAS. WT KRAS protein shuttles between GTP-bound active form and GDP-bound inactive form (left panel). Mut KRAS in the GTP binding domain cannot release GTP, then remained to be constitutively active form of GTP-binding (right panel). KRAS bound GTP induces cell proliferation signal, through the activation of multiple downstream effector pathways, including MAPK and PI3K pathways. Mut KRAS promotes downstream signaling constitutively for tumor cell proliferation and cause uncontrolled cell growth.

RNA interference (RNAi) is a process of post-transcriptional and sequence complementarity-dependent gene silencing triggered by small interfering RNAs (siRNAs), which are double-stranded RNAs with 21 nucleotides in length^12,13^. When siRNA is transfected into cells, it is loaded onto the Argonaute (AGO) protein in the RNA-induced silencing complex (RISC)^14^ and unwound into single-stranded RNAs called guide and passenger strands. The guide strand RNA base-pairs with a target mRNA based on sequence complementarity; then, AGO cleaves the target, suppressing its expression^15,16^. However, it was revealed that a limited fraction of siRNA had been functional in human cells. Therefore, we developed specific sequence rules for designing effective siRNA in mammalian cells^17^. Furthermore, after a series of breakthrough achievements in siRNA therapeutics, patisiran, developed by Alnylam Pharmaceuticals, was approved by the U.S. Food and Drug Administration (FDA) in 2018^18^. To date, five first-generation siRNAs targeting *transthyretin*^19^, *5′-aminolevulinate synthase 1*^20^, *hydroxyacid oxidase 1*^21^, and *proprotein convertase subtilisin / kexin 9*^22^, have been approved by the FDA, and all of these siRNAs satisfied the functional siRNA design rules. These findings demonstrate the unequivocal therapeutic potential of the novel modality. A few siRNAs targeting *KRAS* have been reported^23–25^, but specific siRNAs against each *KRAS* single-nucleotide mutation have not yet been developed. Thus, it is broadly recognized to be a big challenge to selectively suppress Mut *KRAS* without affecting WT *KRAS* expression, as WT *KRAS* is required for the viability and functionality of normal cells^23,26^. Here, we successfully developed a single-nucleotide polymorphism-distinguishable siRNA (SNPD-siRNA) that can discriminate single-nucleotide differences between the WT and Mut *KRAS* genes and suppresses the expression of Mut *KRAS* specifically without affecting the expression of WT *KRAS*.

## RESULTS AND DISCUSSION

### Development of SNPD-siRNA specifically targeting 35G>A mutation in *KRAS* gene

To develop SNPD-siRNA targeting *KRAS*, we first identified the most sensitive position for distinguishing single-nucleotide differences between the siRNA guide strand and its target mRNA using a reporter assay^27^ (Figure S1). The effects on RNAi activity of all possible single-nucleotide mismatches between target mRNA and the siRNA guide strand (positions 1-19) were investigated through quantitative real-time PCR (qRT-PCR). Compared to RNAi activity for a target with a perfectly complementary sequence (black bars in Figure S1), mismatches at the central positions, designated 10 and 11 from the 5′ end of the guide strand, strongly reduced RNAi activity. In addition, considerable effects were observed with mismatches at positions 3–6 in the siRNA seed region.

However, mismatches at other positions, namely 1, 2, 7–9, and 12–19, had no apparent or weak effect. Based on these results, we first constructed SNPD-siKRAS targeting single-nucleotide mutations at nucleotide 35 from the translation initiation site of *KRAS* gene, as 64.2% of KRAS mutations occur at that position (Figure 1B). As G to A mutation (35G>A), corresponding to G12D, has the greatest frequency (34.8%) at nucleotide 35, two reporter plasmids, psiCHECK-KRAS_35G-WT and psiCHECK-KRAS_35A-mut, expressing WT (35G-WT) and 35G>A-mutated (35A-mut) *KRAS* mRNA, respectively, were constructed (Figure S2A).

In order to investigate the effects of central mismatches, siRNA with mismatch at position 9, 10, or 11 to 35G-WT target sequence, but complementary to 35A-mut was firstly synthesized (siKRAS_9m, siKRAS_10m, or siKRAS_11m) (Figure S2B). siKRAS_9m suppressed the RNAi activities against both 35G-WT and 35A-mut targets to similar degrees, siKRAS_10m also suppressed the RNAi activities against both the 35G-WT and 35A-mut targets to similar but weaker degrees.

However, siKRAS_11m showed different levels of RNAi activities on the 35G-WT and 35A-mut targets: the stronger RNAi activity on the 35A-mut target compared to that on the 35G-WT target (Figure S2B and Table S1). However, since its RNAi effect on 35A-mut target was lower compared to that of siKRAS-9m.

To enhance the RNAi effect of siKRAS_11m on 35A-mut target, we secondary changed the terminal nucleotides at both ends, because A/U at the 5’-end of the guide strand and G/C at the 5’-end of the passenger strand is preferable to enhance the RNAi activity^17,28–30^ (Figure S2C). RNAi activities of siKRAS_11m with U at the 5’ end of the guide strand (siKRAS_11m_1U), G at the 5’ end of the passenger strand (siKRAS_11m_19G), and U and G at the 5’ end of the guide and passenger strands, respectively (siKRAS_11m_1U+19G) were examined against both 35G-WT and 35A-mut targets. siKRAS_11m_1U enhanced RNAi activities both on 35G-WT and 35A-mut KRAS targets, but siKRAS_11m_19G suppressed their RNAi activities. However, siKRAS_11m_1U+19G promoted RNAi activity on 35A-mut target to the greatest extent, while its RNAi activity on the 35G-WT target was weak (Figure S2C and Table S1).

Then, thirdly the effects of mismatches at nucleotides 3 to 7 in siKRAS_11m_1U+19G (siKRAS_11m_1U+19G_3m, _4m, _5m, _6m, or _7m) (Figure S2D) were examined, since these positions have secondary strong effects for discriminating single nucleotide differences following those of central region (Figure S1). For 35G-WT targets, siKRAS_11m_1U+19G_3m showed strong RNAi activity, but siKRAS_11m_1U+19G_4m, _5m, _6m, and _7m showed almost no RNAi activities (Figure S2D and Table S1). For 35A-mut targets, siKRAS_11m_1U+19G_3m, _5m, and _6m suppressed the expression of 35A-mut targets at the strong and similar levels, but the effects of siKRAS_11m_1U+19G_4m, and _7m showed weaker effects. These results indicated that siKRAS_11m_1U+19G_5m and _6m are most suitable to distinguish 35G-WT and 35A-mut targets: comparatively strong effects on 35A-mut target but almost no suppression effects on 35G-WT target, although the RNAi effect on 35A-mut target was slightly weak compared to siKRAS-11m_1U+19G.

Then, we fourthly introduced 2’-OMe modifications at positions 6–8 of siKRAS_11m_1U+19G_5m or _6m (siKRAS_11m_1U+19G_5m_OMe6-8, siKRAS_11m_1U+19G_6m_OMe6-8), since these modifications are revealed to enhance RNAi activity^31^. As expected, both of siKRAS_11m_1U+19G_5m_OMe6-8 and siKRAS_11m_1U+19G_6m_OMe6-8 exhibited significantly enhanced RNAi activities on 35A-mut target (Figure S2E and Table S1). Although siKRAS_11m_1U+19G_5m_OMe6-8 showed weak RNAi activity on 35G-WT target, siKRAS_11m_1U+19G_6m_OMe6-8 did not show almost no RNAi activity. Therefore, siRNA with the mismatches at position 6 in addition to position 11 to 35A-mut target and 2’-OMe modifications at positions 6–8 may be most preferable for discriminately suppressing A-mut *KRAS* at position 35 without affecting the expression of WT *KRAS*. We designated the name of this “siKRAS_11m_1U+19G_6m_OMe6-8” (Figure S2E) as “SNPD-siKRAS-35A” (Figure 2A).

**Figure 2.**
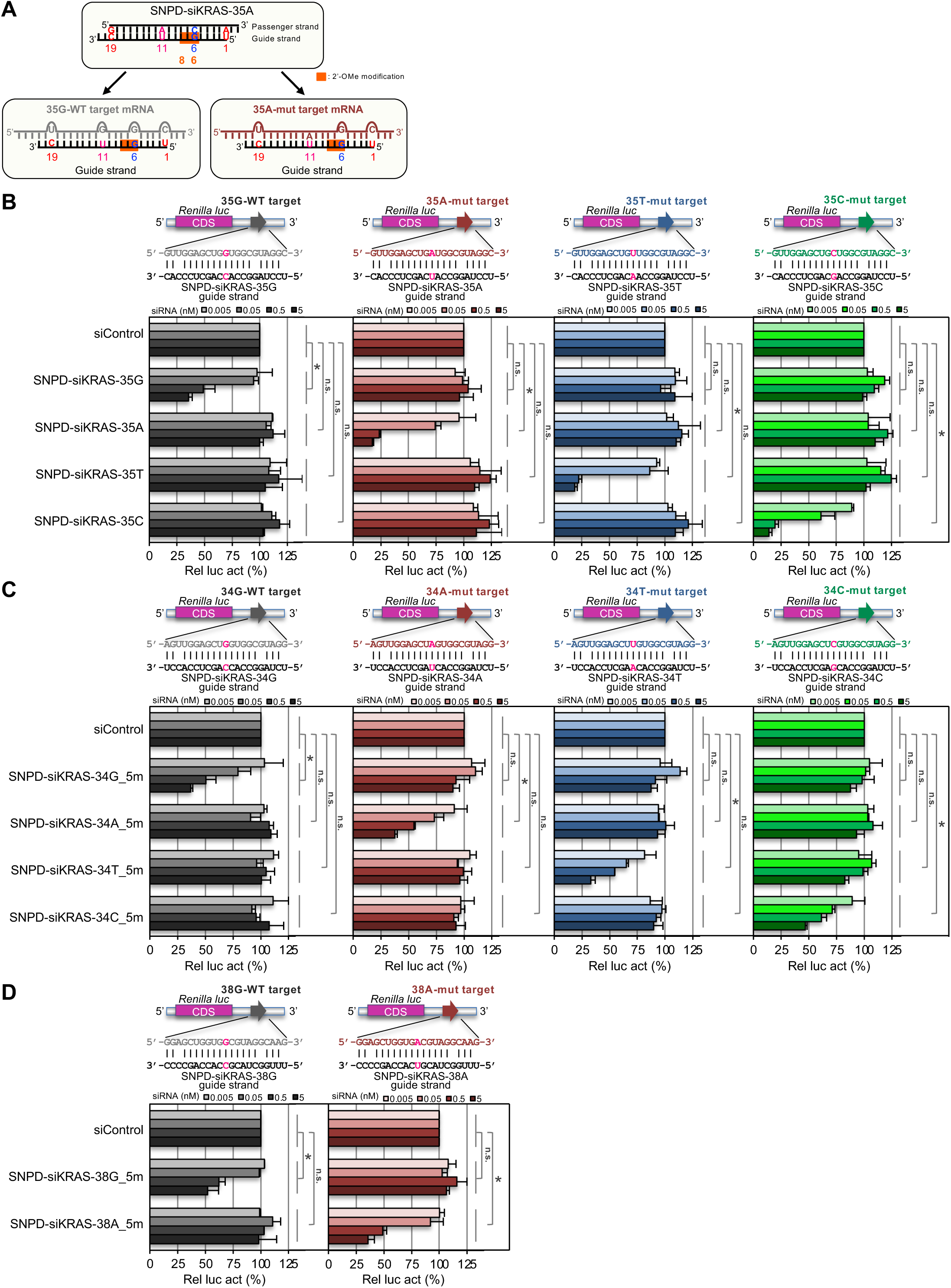
Specific RNAi activity of each SNPD-siKRAS discriminating a single-nucleotide difference. (A) Structure of SNPD-siKRAS-35A and its base-pairing patterns with the targets *KRAS* 35G-WT and 35A-mut. (B) RNAi activities of four types of SNPD-siKRAS-35s against each of the corresponding targets in WT HeLa cells. The psiCHECK-1 plasmid containing the *KRAS* 35G-WT, 35A-mut, 35T-mut, or 35C-mut target sequence in the 3′ UTR of the *Renilla* luciferase gene was used, and the upper panel shows the base-pairing patterns of each SNPD-siKRAS-35 with each target reporter. Dose-dependent RNAi activities of the indicated SNPD-siKRASs against 35G-WT (gray), 35A-mut (dark red), 35T-mut (dark blue), and 35C-mut (green) targets are shown. (C) RNAi activities of four types of SNPD-siKRAS-34s against each of the corresponding targets in WT HeLa cells. The psiCHECK-1 plasmid containing the 34G-WT, 34A-mut, 34T-mut, or 35C-mut target sequence in the 3′ UTR of the Renilla luciferase gene was used, and the upper panel shows the base-pairing patterns of each SNPD-siKRAS-34 with each target reporter. Dose-dependent RNAi activities of the indicated SNPD-siKRASs against 34G-WT (gray), 34A-mut (dark red), 34T-mut (dark blue), and 34C-mut (green) targets are shown. (D) RNAi activities of two types of SNPD-siKRAS-38s against each of the corresponding targets in WT HeLa cells. The psiCHECK-1 plasmid containing the 38G-WT or 38A-mut target sequence in the 3′ UTR of the Renilla luciferase gene was used, and the upper panel shows the base-pairing patterns of each SNPD-siKRAS-38 with each target reporter. Dose-dependent RNAi activities of the indicated SNPD-siKRASs against 38G-WT (gray) and 38A-mut (dark red) targets are shown. All *p*-values were determined using two-way analysis of variance (ANOVA) with reference to the results of siControl (\**p* < 0.05). n.s., not significant.

### Versatility and specificity of SNPD-siKRAS

SNPD-siKRAS-35A has mismatches at positions 6 and 11 with 35G-WT target, but has a mismatch at position 6 alone with 35A-mut, in addition to the mismatches at end positions 1 and 19 for both targets (Figure 2A). Thus, the different mismatched position of SNPD-siKRAS-35A for 35A-mut compared to 35G-WT was only one position 11. This may indicate that SNPD-siKRAS-35A is less likely to base-pair with 35G-WT, since 2 nucleotides mismatches exist at closed sites at positions 6 and 11. However, since SNPD-siKRAS-35A can base-pair with 35A-mut target with one mismatch at position 6, the effective and specific RNAi activity on 35A-mut target may be observed. To ensure the versatility of SNPD-siRNA-35A, other SNPD-siRNAs against other mutations at nucleotide 35 from the translation initiation site of *KRAS* (SNPD-siKRAS-35G (WT), −35T, and −35C) were constructed using the same procedure. Reporter assay showed that SNPD-siKRAS-35A suppressed the 35A-mut target specifically, and no or little RNAi activity against the 35G-WT, 35T-mut, or 35C-mut target was observed (Figure 2B and Table S1). SNPD-siKRAS-35G, −35T, and −35C suppressed the expression of the corresponding target, 35G-WT, 35T-mut, and 35C-mut targets, specifically, with almost no effect on non-targets. Furthermore, we designed SNPD-siKRASs for mutations at nucleotides 34 (codon 12) and 38 (codon 13) (Figure 1A). As mismatches at positions 5 and 6 from the 5′ end of the guide strand showed the similar RNAi activities in the case of SNPD-siKRAS-35A on 35G-WT and 35A-mut targets (Figure S2D), the activities of SNPD-siKRAS-34s and −38s with mismatches at position 5 (5m) or 6 (6m) in addition to position 11, were investigated. SNPD-siKRAS-34G_5m, SNPD-siKRAS-34A_5m, SNPD-siKRAS-34T_5m, SNPD-siKRAS-34C_5m specifically suppressed each of the corresponding targets (Figure 2C and Table S1), whereas SNPD-siKRAS-34G_6m, SNPD-siKRAS-34A_6m, SNPD-siKRAS-34T_6m, SNPD-siKRAS-34C_6m showed negligible effects (Figure S3A and Table S1). Similarly, SNPD-siKRAS-38G_5m and SNPD-siKRAS-38A_5m suppressed each of the corresponding targets (Figure 2D and Table S1), whereas SNPD-siKRAS-38G_6m and SNPD-siKRAS-38A_6m did not (Figure S3B and Table S1). Then, we selected siRNAs with mismatches at position 5 in addition to position 11 in the cases of SNPD-siKRAS-34s and −38s, and designated as SNPD-siKRAS-34G, −34A, −34T, and −34C, and SNPD-siKRAS-38G and −38A, respectively. These results suggest that the strategy used here to design SNPD-siRNA may apply to other target genes, although the mismatch at position 5 or 6 may be dependent on siRNA sequence.

### AGO2 protein is essential for RNAi activity of SNPD-siKRAS

To investigate the mechanism through which SNPD-siRNA distinguishes between WT and Mut *KRAS* mRNAs, the contribution of AGO2 was examined using AGO2 knockout cells generated by the clustered regularly interspaced short palindromic repeats (CRISPR)/Cas9 system^32^ (Figure S4). In AGO2 KO cells, AGO2 mRNA expression was confirmed to be abolished by qRT-PCR (Figure 3A) and AGO2 protein was not detected by western blot (Figure 3B). SNPD-siKRAS-35G, SNPD-siKRAS-35A, SNPD-siKRAS-35T, and SNPD-siKRAS-35C obviously suppressed the expression of each of the corresponding 35G-WT, 35A-mut, 35T-mut, and 35C-mut target in WT HeLa cells (Figure 3C). However, SNPD-siKRAS-35s could not suppress the expression of each of the corresponding targets in AGO2 KO cells (Figure 3C). The lost RNAi activities of SNPD-siKRAS-35s were recovered when AGO2-WT-expression construct was transfected into AGO2 KO cells (Figures 3C and S4E). To confirm these results, we also used the plasmids expressing AGO2 mutants. During RNAi, siRNA loaded on the AGO2 protein is unwound into single-stranded RNAs, and the 5′-terminal of the guide strand RNA is necessary to stably bind to AGO2 in the AGO pocket to cleave the target securely^15,16^. The overexpression of AGO2 mutant which inhibits the binding of the 5’-terminal of the siRNA guide strand^33^ (AGO2-Y529E) and those losing cleavage activities^34^ (AGO2-D597A, AGO2-D669A) led to negligible RNAi activity. Moreover, overexpression of AGO1-WT, AGO3-WT, or AGO4-WT also could not rescue the RNAi activities of SNPD-siKRAS-35s.

**Figure 3.**
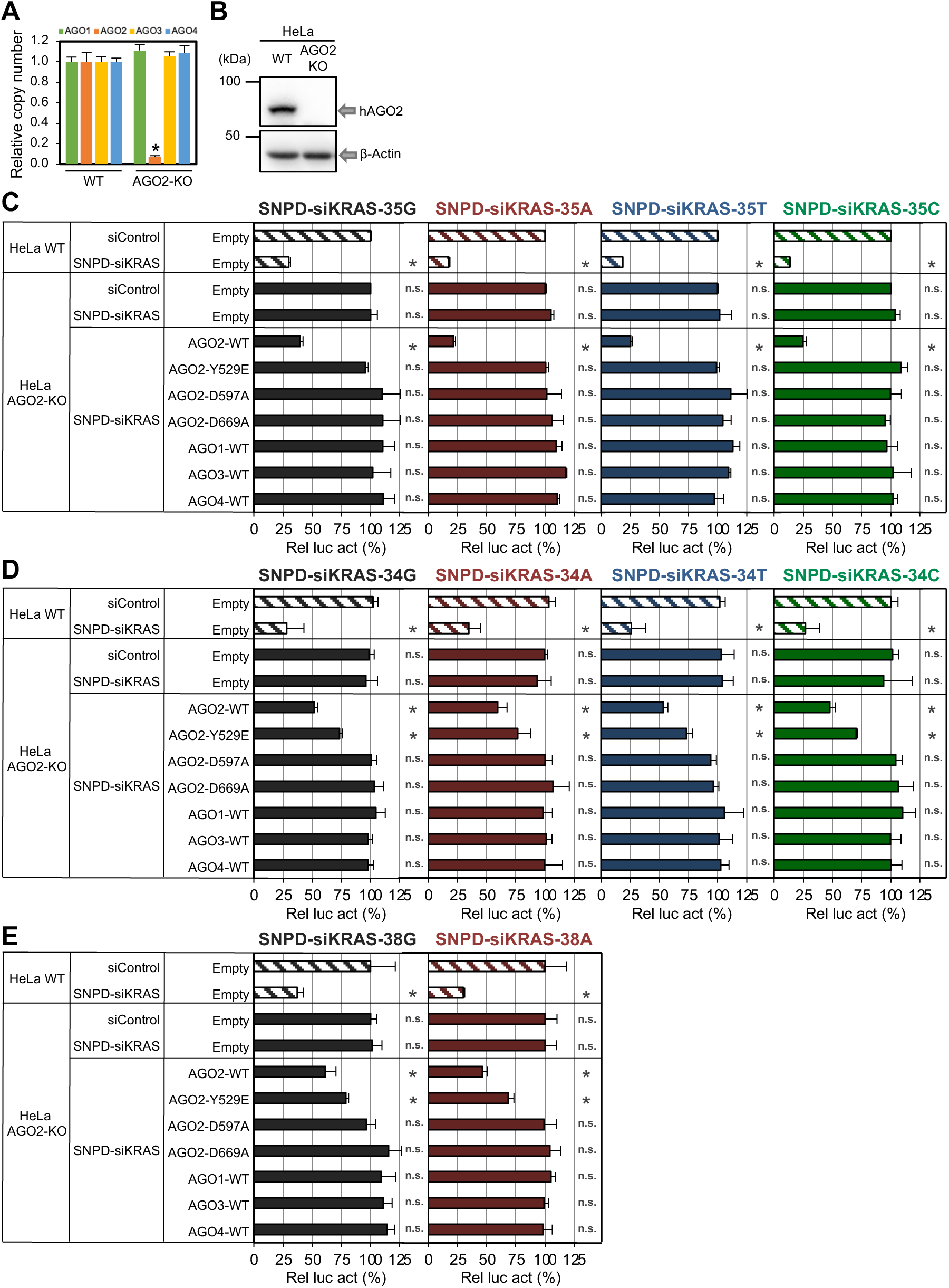
Functional mechanism of SNPD-siKRAS. (A) AGO1-4 mRNA levels in WT and AGO2-KO HeLa cells measured by qRT-PCR. **b.** Western blot of the cell lysate of AGO2-KO HeLa cells detected with anti-AGO2 antibody. (C-E) Rescue experiment of RNAi activities through overexpression of WT AGO2 (pFLAG/HA-AGO2-WT), AGO2 without siRNA binding or cleavage activities (pFLAG/HA-AGO2-Y529E, -D597A, -D669A), and WT AGO1, AGO3, and AGO4 (pFLAG/HA-AGO1-WT, -AGO3-WT, -AGO4-WT) in AGO2-KO cells. SNPD-siKRAS-35s (C), −34s (D) and −38s (E) were transfected at 5 nM. Results are presented as the mean and standard deviation of three independent experiments. All *p*-values were determined using Student′s *t*-test compared with the results obtained in WT HeLa cells transfected with siControl and pFLAG/HA-Empty (\**p* < 0.05). n.s., not significant.

Furthermore, SNPD-siKRAS-34s and −38s also could not repress the expression of each of the corresponding targets in AGO2 KO cells and their RNAi activities were rescued by the overexpression of AGO2-WT but not by AGO2-D597A, AGO2-D669A, AGO1-WT, AGO3-WT, and AGO4-WT (Figures 3D and 3E). Although RNAi activities were only partially recovered when AGO2-Y529E was expressed, it may be consistent with the previous report that small RNAs were only marginally binding with AGO2-Y529E^33^. AGO2 protein is known to cleave target mRNAs at the region between two nucleotides corresponding to the nucleotides 10 and 11 of the siRNA guide strand, whereas other AGO proteins (AGO1, AGO3, and AGO4) have no cleavage activity^34,35^.

SNPD-siKRAS undergoes base-pairing with Mut *KRAS* at the central position 11, but not with WT *KRAS* (Figure 2A), indicating that cleavage of the target by AGO2 at the central position is essential to induce the target-specific RNAi activity of SNPD-siKRAS-35s, −34s, and −38s.

### Validation of RNAi activity of each SNPD-siKRAS on the endogenous *KRAS* expression

The effects of SNPD-siKRAS on endogenous *KRAS* expression were investigated in three human pancreatic cancer-derived cell lines, namely AsPC-1 with homozygous *KRAS* 35G>A mutation, KP-3 with homozygous 35G>T mutation, and PANC-1 with heterozygous 35G>A mutation. Transfection of siControl, SNPD-siKRAS-35G, −35A, or −35T into these cells was performed daily for three consecutive days to introduce siRNA into most of all cells (Figure 4A). One day after the final transfection, *KRAS* mRNA level was measured using qRT-PCR. SNPD-siKRAS-35A decreased the expression of 35A-mut *KRAS* mRNA in AsPC-1 cells, but SNPD-siKRAS-35G and −35T did not, although their mRNA levels were slightly reduced compared to that by siControl (Figure 4B). Similarly, SNPD-siKRAS-35T specifically suppressed the expression of 35T-mut *KRAS* mRNA in KP-3 cells, but SNPD-siKRAS-35G and −35A did not (Figure S5A). These results suggest that SNPD-siKRAS-35A and SNPD-siKRAS-35T could successfully distinguish single-nucleotide differences in endogenous *KRAS* mRNAs. The mutated KRAS is known to continuously activate the downstream MAPK/PI3K pathway, leading to phosphorylation of kinases including extracellular signal-regulated kinase 1/2 (ERK1/2) and thereby promoting tumor cell proliferation^8,9^. Transfection of SNPD-siKRAS-35A, but not SNPD-siKRAS-35G or −35T, reduced the phosphorylation of ERK1/2 in AsPC-1 cells (Figure 4C). Similarly, SNPD-siKRAS-35T, but not SNPD-siKRAS-35G or −35A, reduced the phosphorylated ERK1/2 in KP-3 cells (Figure S5B). To examine the effects of SNPD-siKRAS on cell growth, the number of cells was counted from 1 to 5 days after the final transfection of SNPD-siKRAS. Growth of AsPC-1 and KP-3 cells was significantly suppressed by SNPD-siKRAS-35A and SNPD-siKRAS-35T, respectively, but the growth rates of cells by other SNPD-siKRASs were almost same to that by siControl (Figures 4D and S5C). Furthermore, AsPC-1 cells transfected with SNPD-siKRAS-35A, and KP-3 cells transfected with SNPD-siKRAS-35T were found to be dead in a trypan blue exclusion experiment (Figures 4E and S5D). To investigate whether this cell death was caused specifically by the suppression of Mut *KRAS* but not by WT *KRAS*, the same experiments were performed using PANC-1 cells, which have heterozygous *KRAS* 35G>A mutations. SNPD-siKRAS-35G and SNPD-siKRAS-35A specifically decreased the WT and 35A-mut *KRAS* mRNA levels, respectively (Figure 4F), as determined using specific PCR primers for the WT and 35A-mut (Figure 4G). Furthermore, SNPD-siKRAS-35A reduced phosphorylation of ERK1/2 strongly, but SNPD-siKRAS-35G reduced weakly (Figure 4H). Although both WT and 35A-mut *KRAS* mRNAs were expressed in PANC-1 cells with heterozygous 35G>A mutation, cell growth was repressed by SNPD-siKRAS-35A but not by SNPD-siKRAS-35G (Figure 4I). Also, cell death was specifically induced by SNPD-siKRAS-35A but not by SNPD-siKRAS-35G (Figure 4J). The absolute mRNA level of Mut *KRAS* in PANC-1 cells was approximately 2.5-fold higher than that of WT *KRAS* (Figure 4K), and total *KRAS* mRNA level was approximately 2-fold higher in AsPC-1 cells than in PANC-1 cells (Figure 4L). Thus, since the mRNA levels of Mut *KRAS*s were significantly elevated in both AsPC-1 and PANC-1 cells, cell death may be induced by the absolute downregulation of these *KRAS* mRNA levels. The possibility that Mut *KRAS* has different functions than WT *KRAS* cannot be ruled out.

**Figure 4.**
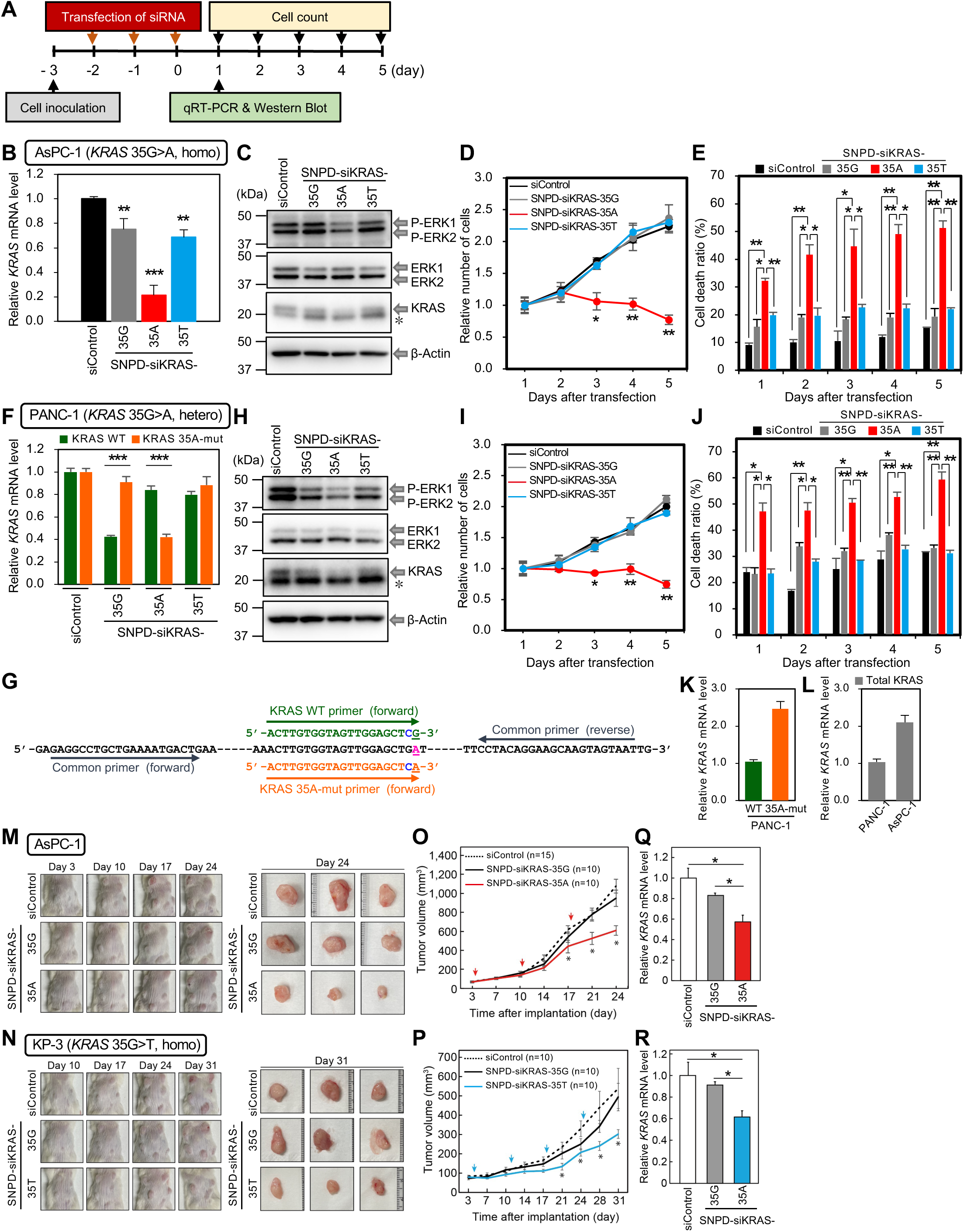
Effects of SNPD-siKRAS on human pancreatic cancer-derived cell lines and a xenograft mouse model. (A) Experimental procedure for investigating the effects of SNPD-siKRAS on pancreatic cancer-derived AsPC-1 and PANC-1 cells. Cells were inoculated at −3 day. Transfection of siRNA was performed at −2, −1, and 0 day. The cells were recovered at 1 day for qRT-PCR and western blot, and cell counts were performed at 1-5 days. (B) *KRAS* mRNA levels were quantified by qRT-PCR using AsPC-1 cells on day 1 after three times transfections of siControl, SNPD-siKRAS-35G, −35A, or −35T at 50 nM. *KRAS* mRNA levels relative to siControl are shown. (C) Western blot of cell lysates of the indicated siRNA-transfected AsPC-1 cells. Unphosphorylated ERK1/2 (ERK1/2) and phosphorylated ERK1/2 (P-ERK1/2) were detected using anti-ERK1/2 and anti-phospho-ERK1/2 antibodies, respectively. KRAS was detected with an anti-Pan-Ras antibody, and β-actin was used as a control. *, non-specific signal. (D) Numbers of AsPC-1 cells relative to the number on day 1 after three times transfections. The black line indicates the number of AsPC-1 cells transfected with siControl, and gray, red, and blue lines indicate the numbers of cells transfected with SNPD-siKRAS-35G, −35A, and −35T, respectively. (E) The ratio of dead AsPC-1 cells on each day. (F) WT and Mut *KRAS* mRNA levels were quantified by qRT-PCR using the specific primer sets shown in (G). (G) PCR primers were used to discriminate WT *KRAS* mRNA from Mut *KRAS* mRNA. Common forward and reverse primers (black) were used to detect both WT and 35A-mut *KRAS* mRNAs simultaneously. The WT and 35A-mut *KRAS* genes were discriminated with specific forward primers designated WT (green) and 35A-mut (orange), respectively. The common primer was used in combination with the WT-specific or 35A-mut KRAS-specific primer. Blue indicates the mismatched nucleotide of each primer. (H) Western blot using lysates of siRNA-transfected PANC-1 cells. Unphosphorylated ERK1/2 (ERK1/2) and phosphorylated ERK1/2 (P-ERK1/2) were detected using anti-ERK1/2 and anti-phospho-ERK1/2 antibodies, respectively. KRAS was detected with an anti-Pan-Ras antibody, and β-actin was used as a control. *, non-specific signal. (I) Numbers of PANC-1 cells relative to the number on day 1 after three times transfections. The black line indicates the number of PANC-1 cells transfected with siControl, and gray, red, and blue lines indicate the numbers of cells transfected with SNPD-siKRAS-35G, −35A, and −35T, respectively. (J) The ratio of dead PANC-1 cells on each day. (K) WT and 35A-mut *KRAS* mRNA levels in PANC-1 cells transfected with siControl. (L) Total *KRAS* mRNA levels in PANC-1 and AsPC-1 cells transfected with siControl. Results are presented as the mean and standard deviation of three independent experiments. All *p*-values were determined using Student′s *t*-test (\**p* < 0.05, \*\**p* < 0.01, \*\*\**p* < 0.001). (M-R) Transfection of SNPD-siKRAS-35A and SNPD-siKRAS-35T significantly inhibited the growth of AsPC-1 (M, O, Q) and KP-3 (N, P, R) cells, respectively, *in vivo*. (M, N) Representative photograph of NOG mice subcutaneously transplanted with AsPC-1 or KP-3 cells (left). Three days after transplantation, tumors had adhered and enlarged. Photograph of tumors extracted from sacrificed mice on the last day of the experiment (right). Each panel shows siControl (top), SNPD-siKRAS-35G (middle), and SNPD-siKRAS-35A or SNPD-siKRAS-35T (bottom) mice. (O, P) Tumor size was measured twice per week from transplantation (day 0). Lines represent the results of siControl (dotted line), SNPD-siKRAS-35G (solid black line), SNPD-siKRAS-35A (solid red line), and SNPD-siKRAS-35T (solid blue line). Red and blue arrows indicate the time of siRNA administration. Results are presented as the mean and standard error of the mean (SEM, n = 10–15). All *p*-values were determined using Student′s *t*-test (\**p* < 0.05). (Q, R) mRNA levels of *KRAS* in tumor tissues extracted from sacrificed mice were evaluated using qRT-PCR. Results are presented as mean and SEM (n = 5). All *p*-values were determined using Student′s *t*-test (\**p* < 0.05).

### Validation of RNAi activity of each SNPD-siKRAS in three-dimensional culture

To mimic the effects of SNPD-siKRAS *in vivo*, we performed three-dimensional spheroid culture of AsPC-1 and KP-3 cells (Figure S6). Sufficiently large cell spheroids were generated after 7 days of three-dimensional culturing, and transfection with each SNPD-siKRAS was then performed daily for three consecutive days (Figure S6A). SNPD-siKRAS-35A significantly reduced the *KRAS* mRNA level to approximately 20% in AsPC-1 cells at one day after the last transfection, and mRNA levels were gradually recovered to the control level about 10 days after transfection (Figure S6B). However, the *KRAS* mRNA levels in the cells transfected with SNPD-siKRAS-35G or −35T were slightly reduced to approximately 70% in AsPC-1 cells at one day after the last transfection, and mRNA levels were recovered at about 7 days (Figure S6B). The growth rate of the AsPC-1 cells transfected with SNPD-siKRAS-35A was slow compared to those transfected with siControl, SNPD-siKRAS-35G, or −35T (Figure S6C). In addition, the absolute sizes of spheroids generated by AsPC-1 cells transfected with SNPD-siKRAS-35A were small compared to those transfected with siControl, SNPD-siKRAS-35G, or −35T (Figure S6D). These results may indicate that the downregulation of *KRAS* mRNA level to approximately 70% in the cells transfected with SNPD-siKRAS-35G or −35T cannot induce almost no effects on the cell growth rate and spheroid sizes (Figures S6C and S6D). However, the *KRAS* mRNA downregulation to about 20% in the cells transfected with SNPD-siKRAS-35A may be enough to repress the cell growth and spheroid sizes. Similar effects of SNPD-siKRAS-35T were observed in KP-3 cells. The *KRAS* mRNA level in KP-3 cells was significantly reduced to approximately 20% at one day after the last transfection of SNPD-siKRAS-35T, and gradually recovered to the control level (Figure S6E). However, the *KRAS* mRNA level by the transfection of siControl, SNPD-siKRAS-35G, or −35A was reduced to about 70% at one day after the last transfection. The growth rate of SNPD-siKRAS-35T-transfected KP-3 cells was slow and their spheroid sizes were small, and almost no effects on the cell growth and spheroid sizes were observed in the cells transfected with siControl, SNPD-siKRAS-35G, or −35A (Figures S6F and S6G). Thus, it was confirmed that SNPD-siKRAS-35A and SNPD-siKRAS-35T exhibit specific effects on each of the corresponding targets in three-dimensional cultures of AsPC-1 and KP-3 cells.

### Effect of SNPD-siKRAS on tumor growth *in vivo* xenograft model

The antitumor function of SNPD-siKRAS-35A or SNPD-siKRAS-35T on pancreatic cancer was further verified using a xenograft *in vivo* mouse model. AsPC-1 or KP-3 cells were subcutaneously implanted into NOG mice. Three days after transplantation, the mice were randomly sorted into three groups and locally treated with 5 µg of a given type of siRNA (siControl, SNPD-siKRAS-35G, SNPD-siKRAS-35A, or SNPD-siKRAS-35T) weekly (Figures 4M and 4N). The average volumes of AsPC-1-derived tumors treated with SNPD-siKRAS-35A were significantly smaller than tumors treated with siControl or SNPD-siKRAS-35G (siControl: 1,076 ± 72 mm^3^, SNPD-siKRAS-35G: 948 ± 88 mm^3^, SNPD-siKRAS-35A: 610 ± 51 mm^3^) (Figure 4O). Similarly, the volumes of KP-3-derived tumors treated with SNPD-siKRAS-35T were significantly reduced (siControl: 539 ± 104 mm^3^, SNPD-siKRAS-35G: 495 ± 71 mm^3^, SNPD-siKRAS-35T: 302 ± 23 mm^3^) (Figure 4P). These findings demonstrate that SNPD-siKRAS-35A and SNPD-siKRAS-35T can inhibit tumor proliferation in an *in vivo* xenograft model. Furthermore, *KRAS* mRNA levels in the recovered implants of AsPC-1 and KP-3 cells were significantly reduced with the administration of SNPD-siKRAS-35A and SNPD-siKRAS-35T, respectively (Figures 4Q and 4R). Thus, the effects of SNPD-siKRAS were confirmed *in vivo*.

## METHODS

### Cell culture

HeLa cells were kindly provided from Prof Hiroyuki Sasaki at Kyushu University. and cultured in Dulbecco’s Modified Eagle’s Medium (FUJIFILM Wako, Osaka, Japan) containing 10% fetal bovine serum (FBS) (Thermo Fisher Scientific, Waltham, MA, USA) and 1% penicillin-streptomycin (PS) solution (FUJIFILM Wako) at 37°C with 5% CO_2_. AsPC-1 (ATCC CRL-1682) and PANC-1 (ATCC CRL-1469) cells were purchased from American Type Culture Collection (Manassas, VA, USA), and KP-3 cells were purchased from Japanese Collection of Research Bioresources Cell Bank (JCRB0178.0, Osaka, Japan). AsPC-1 and KP-3 cells were maintained in RPMI (FUJIFILM Wako) with 10% FBS and 1% PS at 37°C with 5% CO_2_. PANC-1 cells were maintained in DMEM with 10% FBS and 1% PS at 37°C with 5% CO_2_. For 3D spheroid culture, cells were cultured in Dulbecco’s Modified Eagle Medium/Ham F-12 medium (FUJIFILM Wako) with 10% FBS, 1% B-27 supplement, 50 ng/mL EGF, and 50 ng/mL bFGF (Thermo Fisher Scientific) at 37°C with 5% CO_2_.

### Chemical synthesis of siRNA duplexes

RNA oligonucleotides corresponding to the guide and passenger strands of each siRNA were chemically synthesized and subsequently annealed to form siRNA duplexes (Gene Design, Osaka, Japan; Shanghai Gene Pharma, Shanghai, China; SIGMA Inc., Tokyo, Japan). The siRNA sequences used in this study were summarized in Table S2.

### Plasmid construction for reporter assay

Reporter plasmids for measuring RNAi activity of siRNAs were generated using psiCHECK-1 (Promega, Madison, WI, USA). For RNAi activity assays, a 30 nucleotide fragment including WT, G>A, G>T, or G>C mutation at position 35, WT, G>A, G>T, or G>C mutation at position 34, or WT, G>A mutation at position 38 in *KRAS* gene was chemically synthesized (Merck, Darmstadt, Germany) with Xhol or EcoRI sticky end, and the annealed fragment was inserted into the corresponding restriction enzyme sites in the 3’ UTR of *Renilla luciferase* (luc) gene in psiCHECK-1. These constructs were named as following: psiCHECK-KRAS_35G-WT, _35A-mut, _35T-mut, _35C-mut, psiCHECK-KRAS_34G-WT, _34A-mut, _34T-mut, _34C-mut, and psiCHECK-KRAS_38G-WT or _38A-mut. Note that psiCHECK-KRAS_35G-WT, _34G-WT and _38G-WT have identical inserted fragments. The sequences of the inserted fragments were listed in Table S3.

### Measurement of RNAi activity by dual luciferase reporter assay

A cell suspension of HeLa cells (1.0 × 10^5^ cells/mL) was inoculated into a well of 24-well culture plates 1 day before transfection. Cells were simultaneously transfected with 0.005, 0.05, 0.5, or 5 nM of each siRNA, along with 0.1 μg of pGL3-Control vector (Promega), encoding the firefly *luciferase* gene, and 0.01 μg of each psiCHECK construct using Lipofectamine 2000 (Thermo Fisher Scientific). A control siRNA (siControl), which has unrelated KRAS sequences, was used. The transfected cells were lysed with 1×passive lysis buffer (Promega) 24 h after transfection. Luciferase activities were measured using the Dual-Luciferase Reporter Assay System (Promega) and GloMax Discover Microplate Reader (Promega). *Renilla* luciferase activity was normalized by firefly luciferase activity (*Renilla* luciferase activity/firefly luciferase activity), and the relative luciferase activity (Rel luc act) was calculated compared to the result of the control siRNA (siControl) to determine RNA silencing activity.

The half-maximal inhibitory concentration (IC_50_) of siRNA was calculated using the following formula.

IC_50_=10^(LOG_10_(A/B)*(50-C)/(D-C)+LOG_10_(B))

A: Concentration when inhibitory efficiency is lower than 50%

B: Concentration when inhibitory efficiency is higher than 50%

C: Inhibitory efficiency in B

D: Inhibitory efficiency in A

The IC_50_s were shown in Table S1.

### Generation of AGO2-knockout HeLa cells by CRISPR/Cas9 system

Two single-guide RNA (sgRNA), sgAGO2_#1 and sgAGO2_#2, were designed for knockout (KO) of the *AGO2* gene using our online software, CRISPR direct^36^. PX459 vector (Plasmid #62988), expressing SpCas9 and PuromycinR, provided through Addgene (Watertown, MA, USA) was digested by BbsI, and synthetic oligonucleotide (Merck) containing each of two sgRNA fragments was introduced into the BbsI cut site. Two plasmids expressing each of two sgRNAs were co-transfected into HeLa WT cells with Lipofectamine 3000 (Thermo Fisher Scientific). Selection of the transfected cells by puromycin (10 μg/ml) was started on the next day, and single-cell cloning was performed by limiting dilution after 2 days of the selection. The genome DNA sequences of the target site in each of single colonies were checked by direct Sanger sequencing, and the knockout of *AGO2* was confirmed. The sequence primers were listed in Table S4.

### Expression plasmids of AGO proteins

The plasmids expressing AGO1 and AGO2, pIRESneo-FLAG/HA-Ago1 (Plasmid #10820) and - Ago2 (Plasmid #10821), were obtained from Addgene, and FLAG-tagged Ago3 and Ago4 were kindly provided by Dr. Mark A. Kay^37^. These plasmids were called as pFLAG/HA-AGO1-WT, - AGO2-WT, -AGO3-WT, and -AGO4-WT in this study. pFLAG/HA-AGO2-Y529E was constructed in our previous study^38^. For the construction of pFLAG/HA-AGO2-D597A and -D669A, their mutations were introduced into pFLAG/HA-AGO2 by inverse PCR method. The PCR primers were listed in Table S4.

### Transfections into pancreatic cancer-derived cell cultures

For transfection of siRNA into the cultures of pancreatic cancer-derived cells, AsPC-1, KP-3, and PANC-1, cell suspensions (1.0 × 10^5^ cells) were seeded into each well of 24-well culture plate 24 hours before transfection, respectively. These cells were transfected with 50 nM siRNA by Lipofectamine 2000 for 3 consecutive days (Figure 4A). At 24 hours after 3rd transfection, the cells were collected and subjected to cell viability assay, qRT-PCR, and Western blotting. Collected cells were stained with 0.4% (w/v) trypan blue solution (FUJIFILM Wako) and counted cells at 1, 2, 3, 4, and 5 days after the last transfection.

### Transfections in spheroid cultures of pancreatic cancer-derived cells

In the case of transfection in spheroid 3D cultures, AsPC-1 and KP-3 cells cultured for 7 days were collected by centrifugation and resuspended in Opti-MEM medium (Thermo Fisher Scientific). Then siRNA mixed with Lipofectamine RNAiMAX (Thermo Fisher Scientific) in Opti-MEM was added to spheroids. Four hours later, spheroid cultured cells were collected again by centrifugation and re-started spheroid culture in DMEM/Ham F-12 medium with 10% FBS, B-27 supplement, EGF, and bFGF. The photographs of cell morphologies were taken under a microscopy (Keyence, Osaka, Japan).

### Xenograft assay of SNPD-siKRAS

The animal experiments were approved by the animal ethics committee of the University of Tokyo. NOG mice (NOD.Cg-*Prkdc*^scid^*Il2rg*^tm1Sug^/ShiJic, 6 weeks old, male) were obtained from CLEA Japan (Japan) and individually housed in cages with free access to food and water, and reared in 12-hour light-dark cycles in a light-tight chamber at a constant temperature (24 ± 1°C). AsPC-1 and KP-3 cells (2.0 x 10^6^ cells) in Hanks′ Balanced Salt Solution (SIGMA Inc) were implanted into mice subcutaneously. Tumor size was measured by caliper twice a week and the tumor volume was calculated using the following formula: V = L x S^2^ x 1/2 (L: longer diameter of tumor, S: shorter diameter of tumor). Three days after transplantation, 5 µg of each siRNA was injected intratumorally once in a week using *in vivo*-Jet PEI® (Polyplus-transfection Inc., France).

### RNA purification and quantitative qRT-PCR

Total RNA was purified using an Isogen reagent (Nippon gene, Tokyo, Japan). First-strand cDNA was synthesized by High-Capacity cDNA Reverse Transcription kit (Applied Biosystems, Waltham, MA, USA). Quantitative real-time PCR was performed using KAPA SYBR Fast qPCR kit (KAPA Biosystems, Woburn, MA, USA) with QuantiStudio3 real-time PCR system (Applied Biosystems), and analyzed by the ΔΔCt method. The expression level of each sample was normalized by that of glyceraldehyde-3-phosphate dehydrogenase (*GAPDH*). In the case of using a specific primer to discriminately detect WT and 35A-mut *KRAS* genes, their expression levels were analyzed by the absolute quantification method. The PCR primers were listed in Table S4.

### Western blot

The cells were washed with PBS and lysed with 1 × passive lysis buffer (Promega) 24 h after transfection. After determining the protein concentration and denaturation, the samples were separated by sodium dodecyl sulfate-polyacrylamide gel electrophoresis and transferred to an Immobilon-P polyvinylidene fluoride membrane (Merck) using the Mini Trans-Blot Cell (Bio-Rad, Hercules, CA, USA). The membrane was blocked for 1 hour in Tris-buffered saline (TBS) with Tween 20 (TBS-T; 20 mM Tris–HCl [pH 7.5], 150 mM NaCl, 0.1% Tween) supplemented with 5% skim milk (FUJIFILM Wako) at room temperature, and incubated with specific first antibodies in TBS-T with 5% skim milk at 4°C overnight. The membranes were washed three times with TBS-T and reacted with the second antibodies at room temperature for 1 hour. After being washed three times with TBS-T, the membrane was incubated with ECL Prime Western Blotting Detection Reagent (Cytiva, Tokyo, Japan) and visualized using the FUSION (VILBER, Marne-la-Vallée, France). The first antibodies used are as follows; rabbit anti-phospho-ERK1/2 antibody at 1: 1000 dilution (Cell Signaling Technology, Danvers, MA, USA, #4370), rabbit anti-ERK1/2 antibody at 1:1000 dilution (Cell Signaling Technology, #4695), rabbit anti-Pan-RAS antibody at 1: 1000 dilution (Cell Signaling Technology, #3339), mouse anti-FLAG antibody at 1: 1000 dilution (Merck, #F1804), and mouse anti-β-actin antibody at 1: 2000 dilution (Merck, #A2228). The second antibodies used are as follows: horseradish peroxidase (HRP)-linked anti-rabbit or anti-mouse antibody at 1:3000 dilution (Cytiva, #NA934 or #NA931). The protein level of each sample was measured using the software of FUSION, and normalized by that of β-actin.

## SUPPLEMENTAL INFORMATION

Supplemental information can be found online at xxx.

## ACKNOWLEDGEMENTS

We thank Drs. Yukikazu Natori and Kaoru Saigo for the close support, useful discussion, and encouragement throughout the research. K.U.-T. was supported by AMED under Grant Numbers JP22ae0121032 and JP23ae0121031, the START program from the Japan Science and Technology Agency, the GAP fund program from the Division of University Corporate Relations, the University of Tokyo, grant Nos.16K14640 and 26102713 from the Ministry of Education, Culture, Sports, Science and Technology of Japan, and a grant from Japan Health & Research Institute.

## AUTHOR CONTRIBUTIONS

Y. K., H. T., and K. U.-T. designed the study at first, and discussed the methods for experiments. Y. K., A. S., Y. A., and K.U.-T. discussed the way to proceed the research. Y.K. performed the experiments for establishing the framework of SNPD-siRNA and analyzed microarray data, A. S. carried out the experiments using human pancreatic cancer-derived cell lines, and Y. A. implemented the xenograft model experiments. H. T. selected the cell lines for this research. The manuscript was drafted by Y. K., A. S., Y. A., and K.U.-T., and finalized by K.U.-T. All authors read and approved the final manuscript.

---

**Supplementary Figure 1. Reporter assay of RNAi activity depending on single-nucleotide differences.**

RNAi activities of siRNA targeting mRNAs with arbitrary nucleotide sequences were measured using the pTREC system^27^. (A) Structures of pTREC vector and the double-stranded DNA oligonucleotide inserted into the vector. The double-stranded DNA is consisting of two DNA strands of 23-nucleotide-long single-stranded oligonucleotides with *Eco*RI or *Xho*I restriction enzyme site at 5’end, and it was inserted into digested *Eco*RI/*Xho*I site of the vector. PCR primers used to measure the amount of mRNA was shown in the upper region. (B) Target sequences containing all possible single-nucleotide mutations at all possible positions were introduced into the pTREC vector. Each of the resulting vectors was transfected with siRNA into human HeLa cells. The left panel shows the sequences introduced into pTREC vectors: black text indicates the nucleotide matched the siRNA, and the red text indicates a mismatched nucleotide. RNAi activities were determined by measuring mRNA levels transcribed from the pTREC vector via qRT-PCR. The black bar indicates the mRNA level transcribed from the pTREC vector against a perfectly matching target sequence (Extended Data Fig. 1). Arrowhead indicates an mRNA level higher than 50%, representing low RNAi activity.

**Supplementary Figure 2. Development of SNPD-siRNA specifically targeting the 35G>A mutation of the *KRAS* gene.**

(A) Structures of the 35G-WT and 35A-mut reporter constructs (psiCHECK-KRAS_35G-WT and psiCHECK-KRAS_35A-mut, respectively) used for measuring the RNAi activities of siRNAs in WT HeLa cells. The 35G-WT or 35A-mut target region of KRAS was introduced into the 3′ UTR of the *Renilla* luciferase gene. (B–E) **“**siRNA” indicates the name of the siRNA. “siRNA duplex” indicates paired siRNA sequences in which the upper sequence represents the passenger strand sequence from 5′ to 3′ and the lower sequence represents the guide strand sequence from 3′ to 5′. For “Base-pairing pattern with 35G-WT target” and “Base-pairing pattern with 35A-mut target”, each upper sequence indicates the 35G-WT or 35A-mut target sequence transcribed from psiCHECK-KRAS_35G-WT or psiCHECK-KRAS_35A-mut, and the lower sequence indicates the guide strand sequence. siRNA against GFP was used as control siRNA (siControl). Each reporter assay was performed at the indicated concentrations. The horizontal bars indicate relative luciferase activity (Rel luc act). Results are presented as the mean and standard deviation of three independent experiments. (B) Effects of mismatches at positions 9, 10, and 11, respectively. Pink indicates position from the 5’ end of the guide strand complementary with the mutated site of 35A-mut target. (C) Effects of replacement of nucleotides at both ends. Red indicates the substituted nucleotide of siKRAS_11m, and positions 1 or 19 which was substituted with U or C in the guide strand. (D) Effects of additional mismatches in the seed region at positions 3 to 7. Blue indicates nucleotide position mismatched with both 35G-WT and 35A-mut targets. (E) Effects of 2’-OMe modifications at positions 6-8. Orange indicates positions or nucleotides modified with 2’-OMe modification.

**Supplementary Figure 3. RNAi activities of SNPD-siKRASs at nucleotides 34 and 38.**

(A, B) Dose-dependent RNAi activities of the indicated siRNAs against each target in WT HeLa cells. (A) RNAi activities of SNPD-siKRAS-34s with mismatches to the target at positions 6 in addition to position 11. Results for the 35G-WT (gray), 34A-mut (dark red), 34T-mut (dark blue), and 34C-mut (green) targets are shown. (B) RNAi activities of SNPD-siKRAS-38s with mismatches to each target at positions 6 in addition to position 11. Results for the 35G-WT (gray) and 38A-mut (dark red) targets are shown. Relative luciferase activity (Rel luc act) was calculated with reference to the result for siControl to determine RNAi activity. Results are presented as the mean and standard deviation of three independent experiments. All *p*-values were determined using two-way ANOVA with reference to the results of siControl (\**p* < 0.05). n.s., not significant.

**Supplementary Figure 4. Generation of AGO2-KO HeLa cells using the CRISPR/Cas9 system.**

(A) Location of two single guide RNAs used for knockout of AGO2 with the CRISPR/Cas9 system. Magenta indicates the sequence of *AGO2* exon 2. Green and yellow highlights indicate the PAM sequences and sgRNA targeting sites, respectively. Arrowheads indicate the positions of observed double-strand breaks. (B) Genomic DNA sequencing of HeLa AGO2-KO cells. The cleavage site is indicated with a black line. (C) Photographs of WT and AGO2-KO HeLa cells. Bars, 100 µm. (D) Cell growth curve of WT and AGO2 KO HeLa cells. (E) Western blot of AGO2-KO cells and cells transfected with pFLAG/HA-AGO2-WT, -Y529E, -D597A, -D669A, pFLAG/HA-AGO1-WT, - AGO3-WT, or -AGO4-WT against the anti-FLAG antibody. β-actin was used as a control.

**Supplementary Figure 5. Effects of SNPD-siKRAS-35T in KP-3 cell lines with homozygous *KRAS* mutation.**

(A) *KRAS* mRNA levels were quantified using qRT-PCR in KP-3 cells on day 1 after three transfections of siControl, SNPD-siKRAS-35G, −35A, or −35T at 50 nM. (B) Western blot of lysates of siRNA-transfected KP-3 cells. Unphosphorylated ERK1/2 (ERK1/2) and phosphorylated ERK1/2 (P-ERK1/2) were detected with anti-ERK1/2 and anti-phospho-ERK1/2 antibodies, respectively. KRAS was detected with an anti-Pan-Ras antibody. β-actin was used as a control. (C) Numbers of KP-3 cells relative to the number on day 1 after three transfections. The black line indicates the number of KP-3 cells transfected with siControl, and the gray, red, and blue lines indicate the numbers of cells transfected with SNPD-siKRAS-35G, −35A, and −35T, respectively. (D) The ratio of dead KP-3 cells on each day. Results are presented as the mean and standard deviation of three independent experiments. All *p*-values were determined using Student′s *t*-test (\**p* < 0.05, \*\**p* < 0.01, \*\*\**p* < 0.001).

**Supplementary Figure 6. Effects of SNPD-siKRAS on spheroid cultures of pancreatic cancer-derived cells.**

(A) Experimental procedure used to investigate the effects of SNPD-siKRAS on spheroid cultures of AsPC-1 and KP-3 cells. (B) *KRAS* mRNA levels quantified using qRT-PCR in spheroid cultures of AsPC-1 cells on days 1 to 21 after three transfections of siControl, SNPD-siKRAS-35G, −35A, or - 35T at 50 nM relative to the level in siControl on day 1 are shown. (C) Numbers of AsPC-1 cells on days 1 to 21 after three transfections. (D) Photographs of AsPC-1 cells on the indicated day after transfection with each SNPD-siRNA. (E) *KRAS* mRNA levels were quantified using qRT-PCR in the spheroid culture of KP-3 cells on days 1 to 21 after three transfections of siControl, SNPD-siKRAS-35G, −35A, or −35T at 50 nM. (F) Numbers of KP-3 cells on days 1–21 after three transfections. (G) Photographs of KP-3 cells on the indicated day after transfection with each SNPD-siRNA. All *p*-values were determined using Student′s *t*-test (\**p* < 0.05, \*\**p* < 0.01, \*\*\**p* < 0.001).

## REFERENCES

1. Tate, J. G. et al. COSMIC: the Catalogue Of Somatic Mutations In Cancer. Nucleic Acids Res. 47, D941−D947 (2019).

2. Downward, J. Targeting RAS signaling pathways in cancer therapy. Nat. Rev. Cancer 3, 11–22 (2003).

3. Hancock, J. F. Ras proteins: different signals from different locations. Nat. Rev. Mol. Cell Biol. 4, 373–385 (2003).

4. Cox, A. D., Fesik, S. W., Kimmelman, A. C., Luo, J. & Der, C. J. Drugging the undruggable RAS: Mission Possible? Nat. Rev. Drug. Discov. 13, 828–851 (2014).

5. Simanshu, D. K., Nissley, D. V. & McCormick, F. RAS Proteins and Their Regulators in Human Disease. Cell 170, 17–33 (2017).

6. Riely, G. J., Marks, J. & Pao, W. KRAS mutations in non-small cell lung cancer. Proc. Am Thorac. Soc. 6, 201–205 (2009).

7. Fearon, E. R. & Vogelsterin, B. A genetic model for colorectal tumorigenesis. Cell 61, 759–767(1990).

8. Haigis, K. M. KRAS Alleles: The Devil Is in the Detail. Trends Cancer 3, 686–697 (2017).

9. Hunter, J. C. et al. Biochemical and Structural Analysis of Common Cancer-Associated KRAS Mutations. Mol. Cancer, Res. 13, 1325−1335 (2015).

10. Canon, J. et al. The clinical KRAS(G12C) inhibitor AMG 510 drives anti-tumour immunity Nature 575, 217−223 (2019).

11. Hong, D. S. et al. KRAS ^G12C^ Inhibition with Sotorasib in Advanced Solid Tumors. N Engl J Med. 383, 1207−1217 (2020).

12. Fire, A. et al. Potent and specific genetic interference by double-stranded RNA in Caenorhabditis elegans. Nature 391, 806–811 (1998).

13. Hutvagner, G. & Simard, M. J. Argonaute proteins: key players in RNA silencing. Nat. Rev. Mol. Cell Biol. 9, 22–32 (2008).

14. Filipowicz, W. RNAi: The Nuts and Bolts of the RISC Machine. Cell 122, 17–20 (2005).

15. Elbashir, S. M., Martinez, J., Patkaniowska, A., Lendeckel, W. & Tuschl, T. Functional anatomy of siRNAs for mediating efficient RNAi in Drosophila melanogaster embryo lysate. EMBO J. 20, 6877–6888 (2001).

16. Rivas, F. V. et al. Purified Argonaute2 and an siRNA form recombinant human RISC. Nat. Struct. Mol. Biol. 12, 340–349 (2005).

17. Ui-Tei, K. et al. Guidelines for the selection of highly effective siRNA sequences for mammalian and chick RNA interference. Nucleic Acids Res. 32, 936–948 (2004).

18. Adams, D. et al. Patisiran, an RNAi Therapeutic, for Hereditary Transthyretin Amyloidosis. N Engl J Med. 379, 11–21 (2018).

19. Habtemariam, B. A. et al. Single-Dose Pharmacokinetics and Pharmacodynamics of Transthyretin Targeting N-acetylgalactosamine-Small Interfering Ribonucleic Acid Conjugate, Vutrisiran, in Healthy Subjects. Clin Pharmacol Ther. 109, 372–382 (2021).

20. Gonzalez-Aseguinolaza, G. Givosiran-Running RNA Interference to Fight Porphyria Attacks. N Engl J Med. 382, 2366–2367 (2020).

21. Garrelfs, S. F. et al. Lumasiran, an RNAi Therapeutic for Primary Hyperoxaluria Type 1. N Engl J Med. 384, 1216–1226 (2021).

22. Ray, K. K. et al. Inclisiran in Patients at High Cardiovascular Risk with Elevated LDL Cholesterol. N Engl J Med. 376, 1430–1440 (2017).

23. Réjiba, S., Wack, S., Aprahamian, M. & Hajri, A. K-ras oncogene silencing strategy reduces tumor growth and enhances gemcitabine chemotherapy efficacy for pancreatic cancer treatment. Cancer Sci. 98, 1128–1136 (2007).

24. Papke, B. et al. Silencing of Oncogenic KRAS by Mutant-Selective Small Interfering RNA. ACS Pharmacol Transl Sci. 4, 703–712 (2021).

25. Khvalevsky, E. Z. et al. Mutant KRAS is a druggable target for pancreatic cancer. Proc Natl Acad Sci USA 110, 20723–20728 (2013).

26. Koera, K. et al. K-Ras is essential for the development of the mouse embryo. Oncogene 15, 1151–1159 (1997).

27. Ui-Tei, K., Naito, Y. & Saigo, K. Guidelines for the selection of effective short-interfering RNA sequences for functional genomics. Methods Mol. Biol. 361, 201–216 (2007).

28. Khvorova, A., Reynolds, A. & Jayasena, S. D. Functional siRNAs and miRNAs exhibit strand bias. Cell 115, 209–216 (2003).

29. Schwarz, D. S. et al. Asymmetry in the assembly of the RNAi enzyme complex. Cell 115, 199–208 (2003).

30. Noland, C. L. & Doudna, J. A. Multiple sensors ensure guide strand selection in human RNAi pathways. RNA 19, 639–648 (2013).

31. Kobayashi, Y. et al. siRNA Seed Region Is Divided into Two Functionally Different Domains in RNA Interference in Response to 2’-OMe Modifications. ACS Omega 7, 2398–2140 (2022).

32. Jinek, M. et al. A Programmable Dual-RNA–Guided DNA Endonuclease in Adaptive Bacterial Immunity. Science 337, 816–821 (2012).

33. Rüdel, S. et al. Phosphorylation of human Argonaute proteins affects small RNA binding. Nucleic Acids Res. 39, 2330–2343 (2011).

34. Liu, J. et al. Argonaute2 Is the Catalytic Engine of Mammalian RNAi. Science 305, 1437–1441 (2004).

35. Meister, G. et al. Human Argonaute2 Mediates RNA Cleavage Targeted by miRNAs and siRNAs. Mol. cell 15, 185–197 (2004).

36. Naito, Y., Hino, K., Bono, H. & Ui-Tei, K. CRISPRdirect: software for designing CRISPR/Cas guide RNA with reduced off-target sites. Bioinformatics 31, 1120–1123 (2015).

37. Valdmanis, P. N. et al. Expression determinants of mammalian argonaute proteins in mediating gene silencing. Nucleic Acids Res. 40, 3704–3713 (2012).

38. Nishi, K. et al. Control of the localization and function of a miRNA silencing component TNRC6A by Argonaute protein. Nucleic Acids Res. 43, 9856–9873 (2015).

