## Supplemental Figure and Table for "SNPD-siKRAS: siRNA specifically inhibits KRAS with a single-nucleotide mutation"

### Supplementary Figure 1

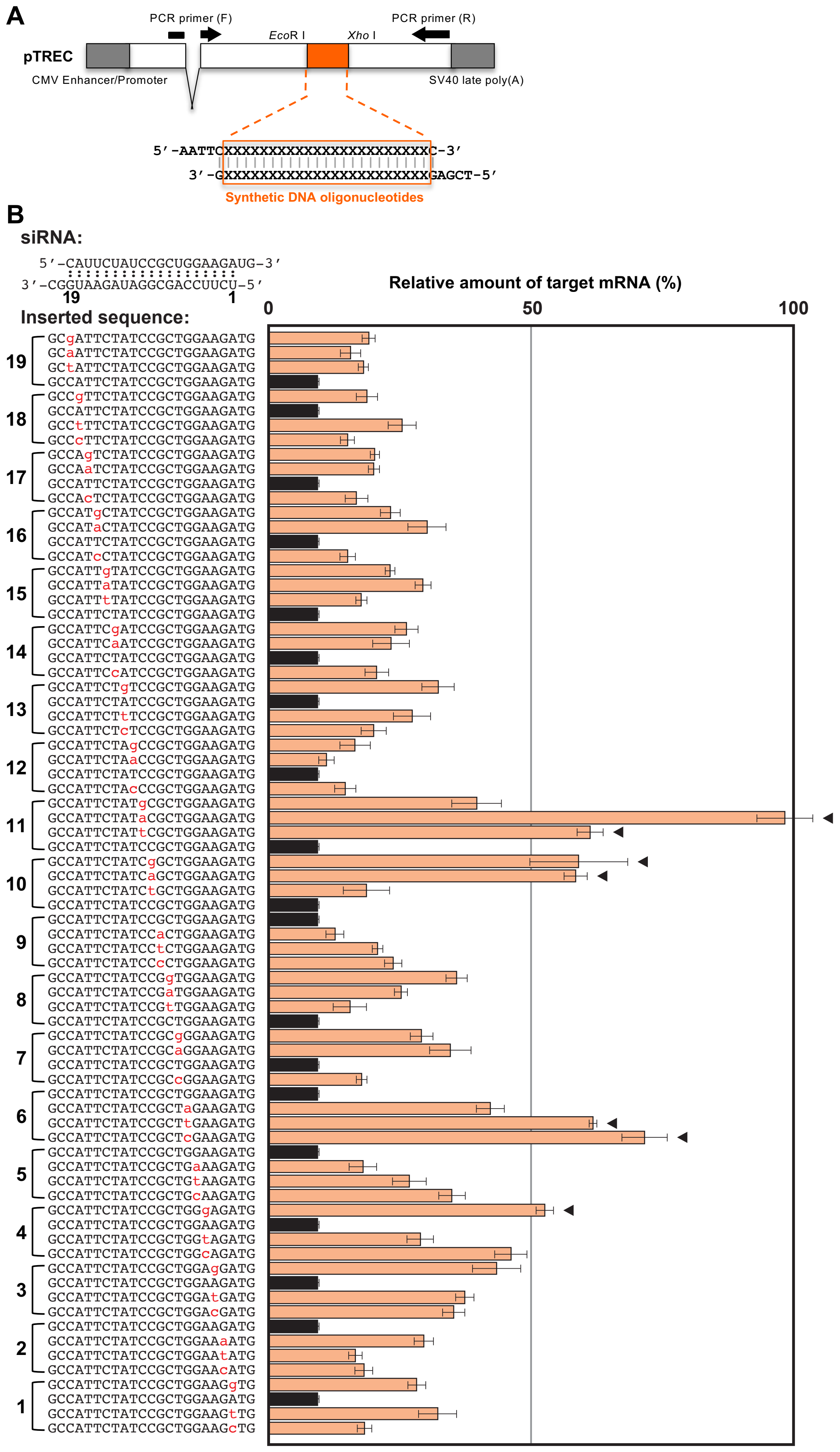

### Supplementary Figure 2

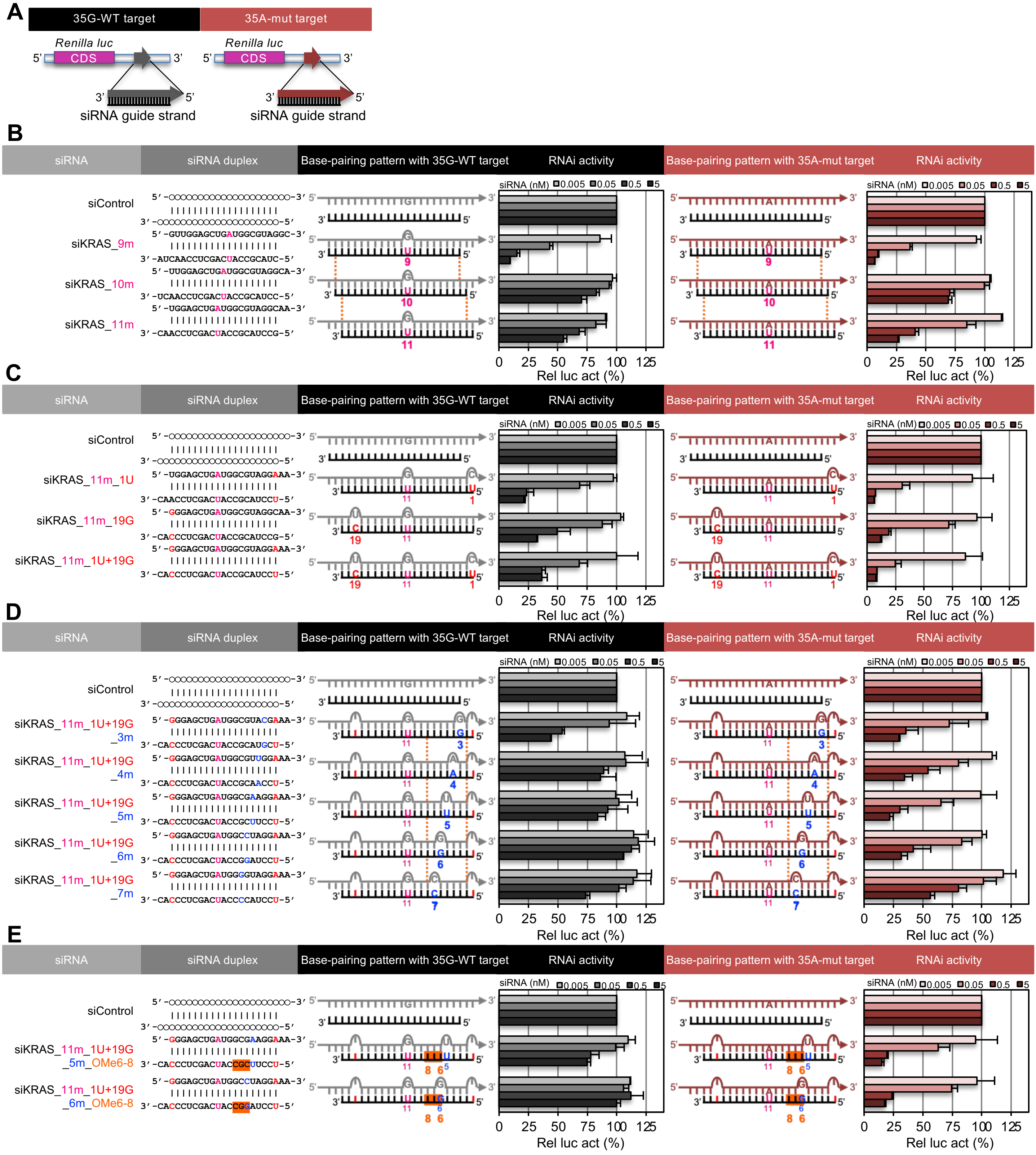

### Supplementary Figure 3

A

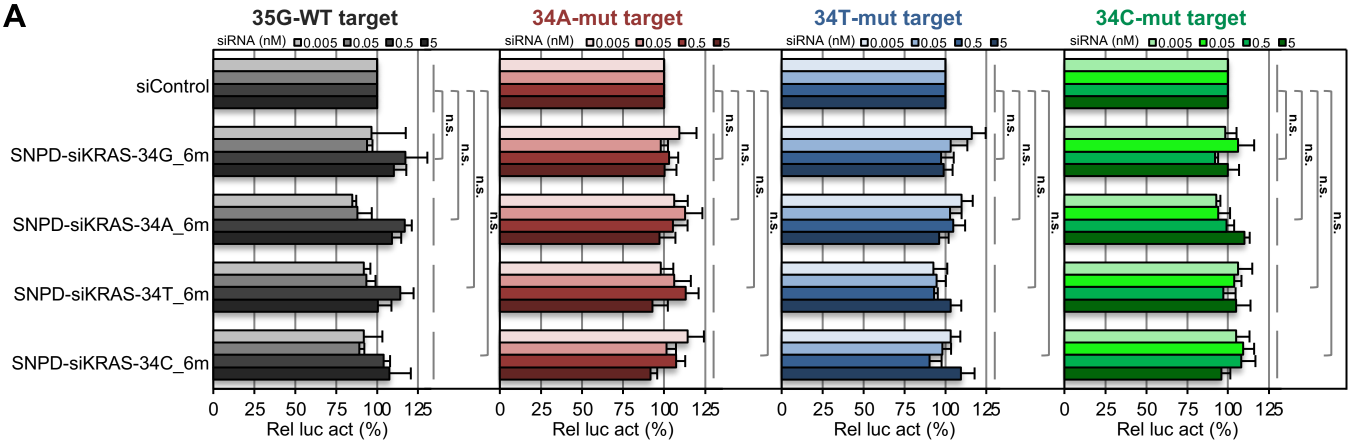


B

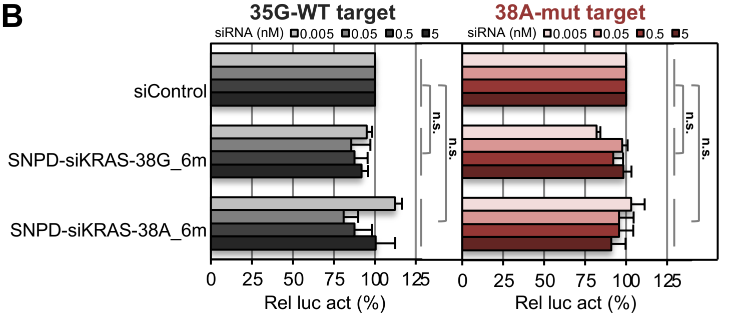

### Supplementary Figure 4

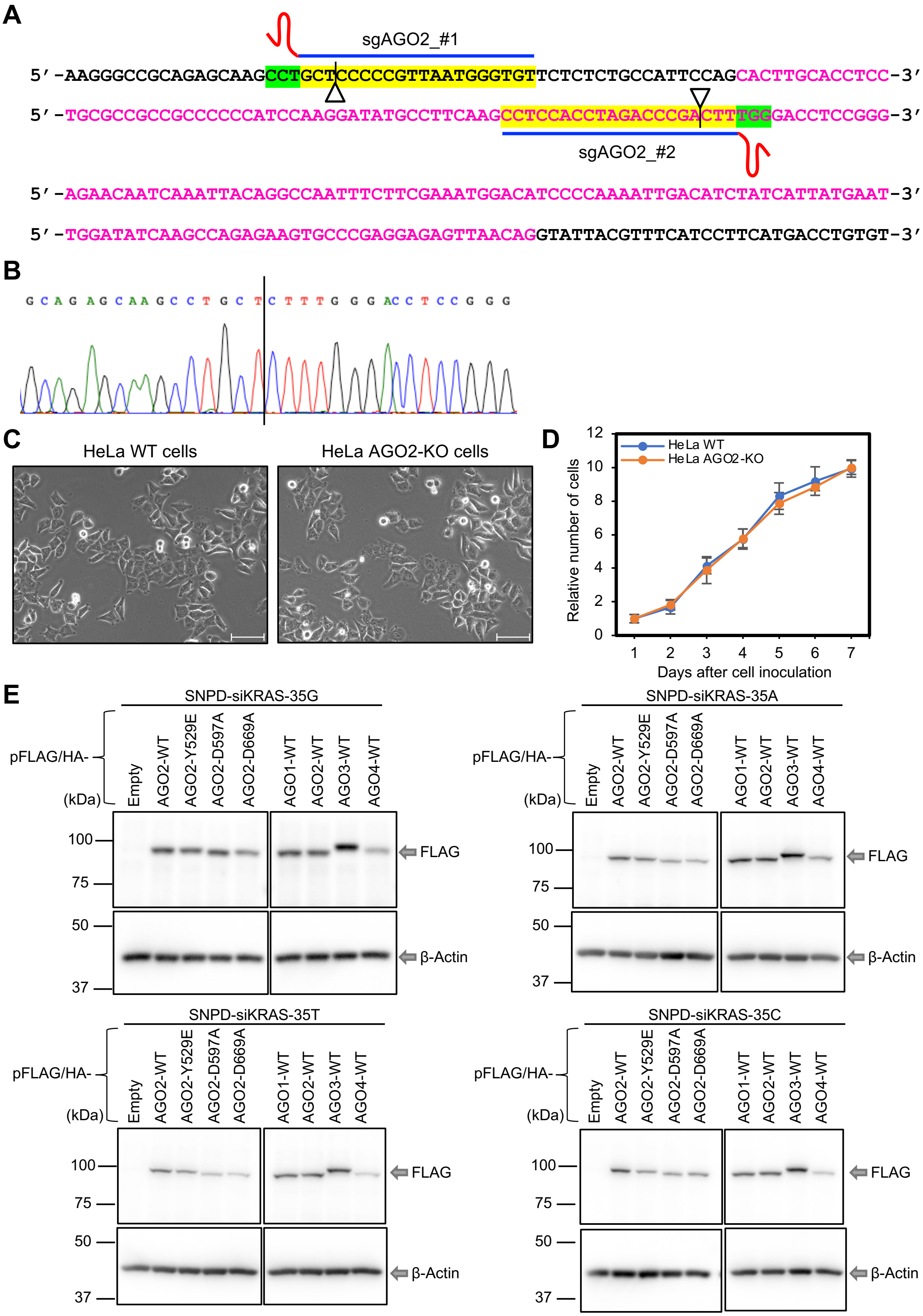

### Supplementary Figure 5

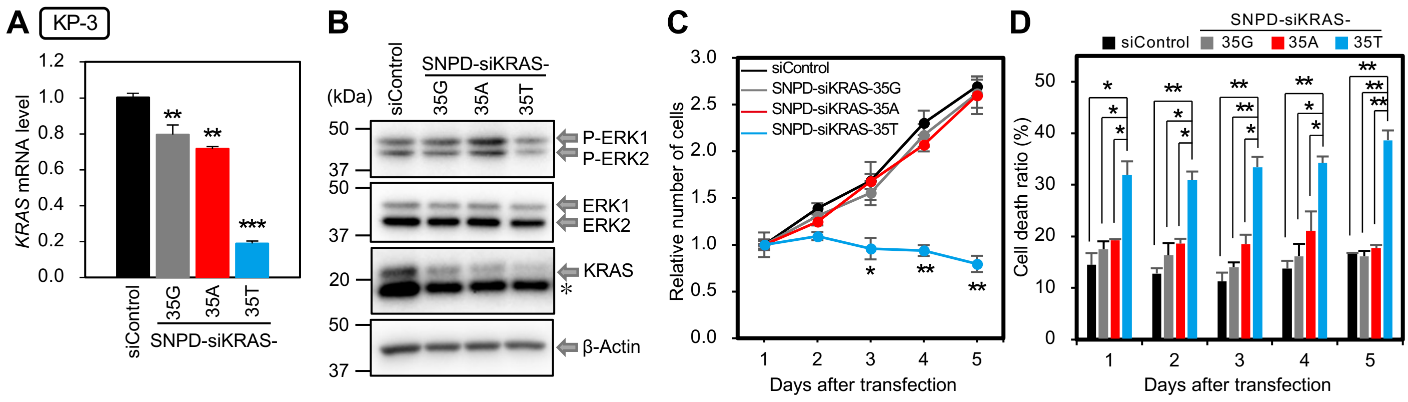

### Supplementary Figure 6

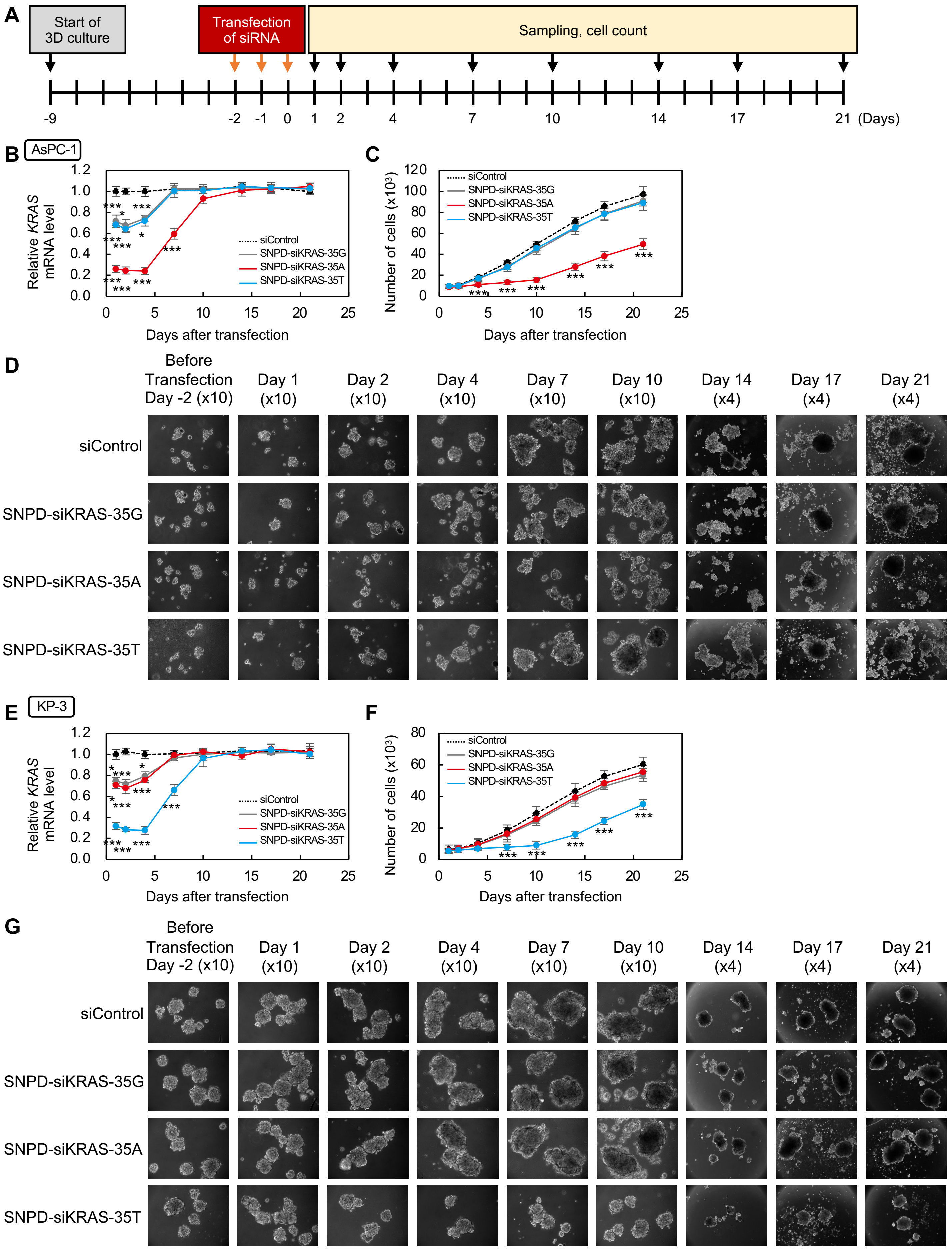

**Supplementary Table 1. IC50 of reporter assay for RNAi activity.**

| siRNA | psiCHECK-KRAS | | | |
|---|---|---|---|---|
| | _35G-WT (pM) | _35A-mut (pM) | _35T-mut (pM) | _35C-mut (pM) |
| siKRAS_9m | 34 | 29 | - | - |
| siKRAS_10m | ∞ | ∞ | - | - |
| siKRAS_11m | ∞ | 311 | - | - |
| siKRAS_11m_1 | 129 | 24 | - | - |
| siKRAS_11m_19 | 480 | 129 | - | - |
| siKRAS_11m_1+19 | 189 | 19 | - | - |
| siKRAS_11m_1+19_3m | 1172 | 202 | - | - |
| siKRAS_11m_1+19_4m | ∞ | 842 | - | - |
| siKRAS_11m_1+19_5m | ∞ | 137 | - | - |
| siKRAS_11m_1+19_6m | ∞ | 315 | - | - |
| siKRAS_11m_1+19_7m | ∞ | ∞ | - | - |
| siKRAS_11m_1+19_5m_OMe6-8 | ∞ | 100 | - | - |
| siKRAS_11m_1+19_6m_OMe6-8 (SNPD-siKRAS-35A) | ∞ | 151 | ∞ | ∞ |
| SNPD-siKRAS-35G | 473 | ∞ | ∞ | ∞ |
| SNPD-siKRAS-35T | ∞ | ∞ | 184 | ∞ |
| SNPD-siKRAS-35C | ∞ | ∞ | ∞ | 93 |

| siRNA | psiCHECK-KRAS | | | |
|---|---|---|---|---|
| | _35G-WT (pM) | _34A-mut (pM) | _34T-mut (pM) | _34C-mut (pM) |
| SNPD-siKRAS-34G_5m | 559 | ∞ | ∞ | ∞ |
| SNPD-siKRAS-34A_5m | ∞ | 932 | ∞ | ∞ |
| SNPD-siKRAS-34T_5m | ∞ | ∞ | 805 | ∞ |
| SNPD-siKRAS-34C_5m | ∞ | ∞ | ∞ | 3081 |
| SNPD-siKRAS-34G_6m | ∞ | ∞ | ∞ | ∞ |
| SNPD-siKRAS-34A_6m | ∞ | ∞ | ∞ | ∞ |
| SNPD-siKRAS-34T_6m | ∞ | ∞ | ∞ | ∞ |
| SNPD-siKRAS-34C_6m | ∞ | ∞ | ∞ | ∞ |

| siRNA | psiCHECK-KRAS | |
|---|---|---|
| | _35G-WT (pM) | _38A-mut (pM) |
| SNPD-siKRAS-38G_5m | ∞ | ∞ |
| SNPD-siKRAS-38A_5m | ∞ | 476 |
| SNPD-siKRAS-38G_6m | ∞ | ∞ |
| SNPD-siKRAS-38A_6m | ∞ | ∞ |

**Supplementary Table 2. SiRNA sequences.**

| siRNA | Passenger strand (5'→3') | Guide strand (5'→3') |
|---|---|---|
| siControl | GCCACAACGUCUAUAUCAUGG | AUGAUAUAGACGUUGUGGCUG |
| siKRAS_9m | GUUGGAGCUGAUGGCGUAGTT | CUACGCCAUCAGCUCCAACTT |
| siKRAS_10m | UUGGAGCUGAUGGCGUAGGCA | CCUACGCCAUCAGCUCCAACU |
| siKRAS_11m | UGGAGCUGAUGGCGUAGGCAA | GCCUACGCCAUCAGCUCCAAC |
| siKRAS_11m_1 | UGGAGCUGAUGGCGUAGGAAA | UCCUACGCCAUCAGCUCCAAC |
| siKRAS_11m_19 | GGGAGCUGAUGGCGUAGGCAA | GCCUACGCCAUCAGCUCCCAC |
| siKRAS_11m_1+19 | GGGAGCUGAUGGCGUAGGAAA | UCCUACGCCAUCAGCUCCCAC |
| siKRAS_11m_1+19_3m | GGGAGCUGAUGGCGUACGAAA | UCGUACGCCAUCAGCUCCCAC |
| siKRAS_11m_1+19_4m | GGGAGCUGAUGGCGUUGGAAA | UCCAACGCCAUCAGCUCCCAC |
| siKRAS_11m_1+19_5m | GGGAGCUGAUGGCGAAGGAAA | UCCUUCGCCAUCAGCUCCCAC |
| siKRAS_11m_1+19_6m | GGGAGCUGAUGGCCUAGGAAA | UCCUAGGCCAUCAGCUCCCAC |
| siKRAS_11m_1+19_7m | GGGAGCUGAUGGGGUAGGAAA | UCCUACCCCAUCAGCUCCCAC |
| siKRAS_11m_1+19_5m_OMe6-8 | GGGAGCUGAUGGCGAAGGAAA | UCCUU<u>CGC</u>CAUCAGCUCCCAC |
| siKRAS_11m_1+19_6m_OMe6-8 (SNPD-siKRAS-35A) | GGGAGCUGAUGGCCUAGGAAA | UCCUA<u>GGC</u>CAUCAGCUCCCAC |
| SNPD-siKRAS-35G | GGGAGCUGGUGGCCUAGGAAA | UCCUA<u>GGC</u>CACCAGCUCCCAC |
| SNPD-siKRAS-35T | GGGAGCUGUUGGCCUAGGAAA | UCCUA<u>GGC</u>CAACAGCUCCCAC |
| SNPD-siKRAS-35C | GGGAGCUGCUGGCCUAGGAAA | UCCUA<u>GGC</u>CAGCAGCUCCCAC |
| SNPD-siKRAS-34G_5m | GUGGAGCUGGUGGCCUAGACA | UCUAG<u>GCC</u>ACCAGCUCCACCU |
| SNPD-siKRAS-34A_5m | GUGGAGCUAGUGGCCUAGACA | UCUAG<u>GCC</u>ACUAGCUCCACCU |
| SNPD-siKRAS-34T_5m | GUGGAGCUUGUGGCCUAGACA | UCUAG<u>GCC</u>ACAAGCUCCACCU |
| SNPD-siKRAS-34C_5m | GUGGAGCUCGUGGCCUAGACA | UCUAG<u>GCC</u>ACGAGCUCCACCU |
| SNPD-siKRAS-34G_6m | GUGGAGCUGGUGGGGUAGACA | UCUAC<u>CCC</u>ACCAGCUCCACCU |
| SNPD-siKRAS-34A_6m | GUGGAGCUAGUGGGGUAGACA | UCUAC<u>CCC</u>ACUAGCUCCACCU |
| SNPD-siKRAS-34T_6m | GUGGAGCUUGUGGGGUAGACA | UCUAC<u>CCC</u>ACAAGCUCCACCU |
| SNPD-siKRAS-34C_6m | GUGGAGCUCGUGGGGUAGACA | UCUAC<u>CCC</u>ACGAGCUCCACCU |
| SNPD-siKRAS-38G_5m | GGCUGGUGGCGUAGCCAAAAG | UUUGG<u>CUA</u>CGCCACCAGCCCC |
| SNPD-siKRAS-38A_5m | GGCUGGUGACGUAGCCAAAAG | UUUGG<u>CUA</u>CGUCACCAGCCCC |
| SNPD-siKRAS-38G_6m | GGCUGGUGGCGUACGCAAAAG | UUUGC<u>GUA</u>CGCCACCAGCCCC |
| SNPD-siKRAS-38A_6m | GGCUGGUGACGUACGCAAAAG | UUUGC<u>GUA</u>CGUCACCAGCCCC |

underline = 2'-OMe modification, KRAS = Kirsten rat sarcoma viral oncogene homolog

**Supplementary Table 3. Inserted oligonucleotide sequences in psiCHECK-reporters.**

| Oligonucleotide name | Sequence (5'→3') |
|---|---|
| KRAS_35G-WT_s | tcgagTGGTAGTTGGAGCTGGTGGCGTAGGCAAGAGTGg |
| KRAS_35G-WT_as | aattcCACTCTTGCCTACGCCACCAGCTCCAACTACCAc |
| KRAS_35A-mut_s | tcgagTGGTAGTTGGAGCTGATGGCGTAGGCAAGAGTGg |
| KRAS_35A-mut_as | aattcCACTCTTGCCTACGCCATCAGCTCCAACTACCAc |
| KRAS_35T-mut_s | tcgagTGGTAGTTGGAGCTGTTGGCGTAGGCAAGAGTGg |
| KRAS_35T-mut_as | aattcCACTCTTGCCTACGCCAACAGCTCCAACTACCAc |
| KRAS_35C-mut_s | tcgagTGGTAGTTGGAGCTGCTGGCGTAGGCAAGAGTGg |
| KRAS_35C-mut_as | aattcCACTCTTGCCTACGCCAGCAGCTCCAACTACCAc |
| KRAS_34A-mut_s | tcgagTGGTAGTTGGAGCTAGTGGCGTAGGCAAGAGTGg |
| KRAS_34A-mut_as | aattcCACTCTTGCCTACGCCACTAGCTCCAACTACCAc |
| KRAS_34T-mut_s | tcgagTGGTAGTTGGAGCTTGTGGCGTAGGCAAGAGTGg |
| KRAS_34T-mut_as | aattcCACTCTTGCCTACGCCACAAGCTCCAACTACCAc |
| KRAS_34C-mut_s | tcgagTGGTAGTTGGAGCTCGTGGCGTAGGCAAGAGTGg |
| KRAS_34C-mut_as | aattcCACTCTTGCCTACGCCACGAGCTCCAACTACCAc |
| KRAS_38A-mut_s | tcgagTGGTAGTTGGAGCTGGTGACGTAGGCAAGAGTGg |
| KRAS_38A-mut_as | aattcCACTCTTGCCTACGTCACCAGCTCCAACTACCAc |

_s = sense strand; _as = antisense strand, lower case = sequence of restriction enzyme site

**Supplementary Table 4. Primer sequences.**

| Primer name | Sequence (5'→3') |
|---|---|
| AGO2-KO-sequence_F | AATGGAAACAGCAGAGGGGG |
| AGO2-KO-sequence_R | CGCAGACCACTTACACAGGT |
| AGO2-D597A_mutagenesis_F | GCCGTCACTCACCCCCCCGCCGG |
| AGO2-D597A_mutagenesis_R | TGCTCCCAGAAAGATGACGGGCTGCTGGAAC |
| AGO2-D669A_mutagenesis_F | GCCGGTGTCTCTGAAGGCCAGTTCCAGC |
| AGO2-D669A_mutagenesis_R | GCGGTAGAAGATGATGCGGGTGGG |
| KRAS common_F | GAGGCCTGCTGAAAATGACTG |
| KRAS common_R | ATTACTACTTGCTTCCTGTAGG |
| KRAS WT_F | ACTTGTGGTAGTTGGAGCTCG |
| KRAS 35A-mut_F | ACTTGTGGTAGTTGGAGCTCA |
| GAPDH_F | TGCACCACCAACTGCTTAG |
| GAPDH_R | AGAGGCAGGGATGATGTTC |

_F = forward primer; _R = reverse primer
